# Analyzing engineered point spread functions using phasor-based single-molecule localization microscopy

**DOI:** 10.1101/2020.04.15.043182

**Authors:** Koen J.A. Martens, Abbas Jabermoradi, Suyeon Yang, Johannes Hohlbein

## Abstract

The point spread function (PSF) of single molecule emitters can be engineered in the Fourier plane to encode three-dimensional localization information, creating double-helix, saddle-point or tetra-pod PSFs. Here, we describe and assess adaptations of the phasor-based single-molecule localization microscopy (pSMLM) algorithm to localize single molecules using these PSFs with sub-pixel accuracy. For double-helix, pSMLM identifies the two individual lobes and uses their relative rotation for obtaining *z*-resolved localizations, while for saddle-point or tetra-pod, a novel phasor-based deconvolution approach is used. The pSMLM software package delivers similar precision and recall rates to the best-in-class software package (SMAP) at signal-to-noise ratios typical for organic fluorophores. pSMLM substantially improves the localization rate by a factor of 2 - 4x on a standard CPU, with 1-1.5·10^4^ (double-helix) or 2.5·10^5^ (saddle-point/tetra-pod) localizations/second.

## 1. Introduction

Fluorescence microscopy is frequently employed in biological sciences due to its high selectivity and non-invasiveness. Conventionally, the obtainable optical resolution in fluorescence microscopy is given by Abbe’s diffraction limit which is equal to the wavelength of the light divided by double the numerical aperture of the objective (∼ 200 nm for visible light). A multitude of techniques summarized by the term super-resolution (SR) microscopy or nanoscopy [1–3] have been developed, however, to obtain spatial information well below this limit. These techniques include (d)STORM (direct stochastic optical reconstruction microscopy) [4,5], PALM (photo-activatable localization microscopy) [6], SIM (structured illumination microscopy) [7], STED (stimulated emission depletion microscopy) [8], RESOLFT (reversible saturable optical fluorescence transitions) [9], SOFI (super-resolution optical fluctuation imaging) [10], SRRF (super-resolution radial fluctuations) [11] and MINFLUX (minimal photon fluxes localization microscopy) [12].

Single-molecule localization microscopy (SMLM) is the sub-collection of super-resolution techniques in which the fluorescent emission profile, ordinarily referred to as a point spread function (PSF), of a single fluorophore is localized with a precision (∼ 5 - 40 nm) that can exceed the classical resolution limit by more than one order of magnitude [13–16]. SMLM is therefore an integral part of STORM and PALM, and has been extensively used in biological research [17–19], for example to study DNA transcription [20,21], CRISPR-Cas DNA screening [22–24], nuclear pore complexes [25,26], and microtubules [27].

In a conventional fluorescence microscope, a PSF from a single emitter in focus resembles an Airy pattern, which can be approximated by a 2-dimensional Gaussian function. This approach has been the basis of the earliest localization algorithms [13,16,28], which allow for determination of the emitter locations [16] as long as overlapping of PSFs is negligible. Besides Gaussian-based methods, these symmetrical PSFs have been analyzed via other mathematical frameworks, such as radial symmetry [29], cubic splines [30], or phasor (Fourier) analysis [31].

The shape of the PSF quickly deteriorates, however, if the emitter is out of focus (∼100s of nm), leading to both a limited available axial range and inaccessibility of the absolute axial position [32]. Therefore, a variety of methods have been developed to modulate the shape of the PSF depending on the emitter’s axial position [33]. Historically, the first method (astigmatism; AS) introduced a cylindrical lens in the emission pathway to create ellipsoid PSFs if the emitters are out of focus [34,35]. The extent of the deformation along with its orientation allows for determination of the axial position after a calibration procedure, and fitting of these PSFs could usually be performed by derivatized localization algorithms as the ones used for 2D PSFs [28,31,36]. However, the available axial range of astigmatism is limited to less than ∼1 µm, which lead to the development of more advanced PSF shaping procedures that involve modulating the light in the pupil (Fourier) plane. Using a spatial light modulator (SLM), the principle was first employed to create a double helix (DH) pattern, in which the PSF is split in two separate lobes that non-degeneratively rotate around each other based on the emitters axial position, resulting in an usable axial range up to 2.5 µm [37]. Later, the same group theoretically maximized the information content of PSFs resulting in the Saddle-Point (SP) or Tetra-Pod (TP) designs, which are suitable for 3 µm (SP) or ≥ 6 µm (TP) axial ranges [38,39]. PSFs for both SP and TP are altered in the Fourier plane via a phase mask [38,39] or deformable mirror [40].

Determining the sub-pixel positions corresponding to the emitters via DH, SP, or TP PSFs, however, is more challenging than for symmetric or AS PSFs, as fitting with a single 2D Gaussian is insufficient. The current state-of-the-art fitting algorithms [41] rely on phase retrieving methods [40,42] or spline interpolation [26] to determine a PSF model based on calibration samples. A high-resolution PSF model can then be determined from these models which is fitted on experimental data. These methodologies can work with arbitrarily shaped PSFs, including DH, SP and TP. However, these methods are computationally expensive and thus time-consuming. Recently, real-time fitting localization of experimental PSFs have been achieved using graphical processing units (GPUs) [26], but this has not yet been achieved on computation processing units (CPUs), which would increase the accessibility and might allow implementations directly on the camera hardware.

Here, we show fast retrieval of DH (1.5·10^4^ loc/s) and SP/TP (2.5·10^5^ loc/s) PSF localizations on a standard CPU via novel adaptations of the phasor-based single-molecule localization microscopy (pSMLM) algorithm [31]. We first explain the underlying methodology for DH and for SP/TP, termed circular-tangent (ct-)pSMLM, and then explore the performance of the methods by analyzing simulated and experimental data. We have implemented all pSMLM versions (2D, AS, DH, SP/TP) in a recently published software package (SMALL-LABS [43]), resulting in user-friendly and open-source software to quickly perform sub-pixel localization including advanced background filtering options.

## 2. Material and methods

### 2.1 Software and hardware

All software was written and ran in MATLAB (MathWorks, UK) version 2018b on a 64-bit Windows 10 computer equipped with an Intel i5-8600 CPU @ 3.10 GHz, 16 GB RAM.

### 2.2 SMALL-LABS software

Our software package expands the original SMALL-LABS software [43] in several ways. Firstly, we added the original pSMLM-3D algorithm for 2D or astigmatism PSF sub-pixel localization, as well as the novel variations discussed in this manuscript. Next, a custom GUI was written to increase user accessibility. Lastly, the pre- and post-processing options are expanded with wavelet filtering [44], cross-correlation drift correction in three dimensions [45], and average shifted histogram result image generation [46,47].

### 2.3 Saddle-point PSF simulations

PSF simulations have been performed as described earlier [16,31] with NA = 1.25, emission light at 500 nm, 100 nm/pixel camera acquisition and 1000 PSFs for every intensity/noise combination. We used a full vectorial model of the PSF needed to describe the high NA case typically used in fluorescent super-resolution imaging. The center of the PSF is located within ± 1 pixel of the center of the image. Zernike polynomials 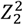 (primary astigmatism) and 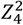 (secondary astigmatism) are introduced in a 0.5:-0.65 ratio [40], and *z*-positions were chosen randomly between –1.5 and +1.5 µm away from the focal plane.

### 2.4 Sub-pixel localization of single-molecule data

For double-helix (DH) sub-pixel localization, the four datasets from the 2016 SMLM challenge [41] were analyzed, which use experimental PSF models. These datasets differ in signal-to-noise (SNR) values (‘high SNR’, which mimics Alexa647 fluorophores, and ‘low SNR’, which mimics fluorescent proteins) and in emitter density (‘low density’ (LD) at 0.2 loc/µm^2^ and ‘high density’ (HD) at 2 loc/µm^2^).

For double-helix localization, the following settings were used. For SMALL-LABS-pSMLM-DH, the temporal window length and the minimum duration of fluorophore on-time before it is discarded were both set to 150 frames. Filtering for region-of-interest (ROI) finding was performed with a β-Spline wavelet filter with the threshold set to 1.9 times the standard deviation of a filtered frame. Single-lobe DH location was performed with a phasor radius of 4 pixels (low density) or 2 pixels (high density). The *z*-position was calculated via a calibration with identical phasor radius. For the SMAP software with fit3dSpline sub-pixel localization [26], a calibration was performed with a 33×33 pixel ROI. Then, localizations were identified via a mean calibrated PSF, with a 2.9 pixel Gaussian blur, using all calibrated *z*-positions. A threshold set to an absolute cutoff value of 86 (high SNR), 76 (low SNR, LD), or 29 (low SNR, HD) photons was used. The calibrated spline PSF was fitted with a 15 × 15 pixel ROI. Then, localizations with relative log-likelihood lower than - 2 (low density) or −5 (high density) were discarded.

For the localization of simulated saddle-point PSFs, we note that in order to prevent localization artefacts at specific *z* positions for ct-pSMLM (SI fig 1), a large ROI (> ∼ 1.2x the maximum distance between the lobes) had to be used to determine lateral localization accurately, while a smaller ROI (∼ 2.3 × 2.3 µm) was required for accurate axial localization, as ct-pSMLM with a large ROI cannot accurately describe the axial position around the focus. Then, all PSFs were localized directly with ct-pSMLM as described in the result section or via the SMAP software with fit3dSpline sub-pixel localization [26]. For ct-pSMLM, we used a 23×23 pixel ROI to calculate the *z*-position and a 43 × 43 pixel ROI to calculate the *x* and *y* position, and a 4^th^-order polynomial was used to fit the calibration curve. For SMAP, calibration was performed with a 15-px Gaussian blur to find PSFs, a 51 × 51 pixel ROI, and 10 nm axial distance between every localization. For localization, a 10-px Gaussian blur was used to find PSFs, with a threshold of 20. A 41 × 41 pixel ROI spline fitting with 100 iterations based on the calibrated data was used to localize the PSFs.

Localization of experimental saddle-point data was performed with the SMALL-LABS-pSMLM software package. A median background subtraction with temporal window length of 150 frames and minimum duration of fluorophores to be discarded of 100 frames was used. Localizations were identified via a bandpass filter with a 95 threshold percentile. Potential lobes of saddle-point point spread functions were identified with a 3 pixel radius ROI 2D phasor fitting routine. Ct-pSMLM fitting was then performed with an 11 pixel radius ROI around the center of localizations. Calibration was performed using simulated point spread functions at varying *z* positions, consisting of 5000 photons on a noiseless background, with deformations similar to experimental data. Three-dimensional cross correlative drift correction was performed via the SMALL-LABS-JH software, with 10 lateral subpixels and 10 temporal bins. The average shifted histogram image was created using ThunderSTORM [46], using 50 nm axial bins and 10 lateral subpixels.

### 2.5 Assessment of localization performance

For double-helix (DH), localizations between ground-truth (GT) and software 1 (S1; SMALL-LABS-pSMLM-DH) and between GT and software 2 (S2; SMAP with fit3dSpline) are linked on a frame-by-frame basis, with a maximum allowed lateral distance of 250 nm, and a maximum allowed axial distance of 500 nm. The median offset between GT and S1 and between GT and S2 is calculated and subtracted from the S1 and S2 datasets, to avoid introducing consistent offset errors in the RMSE calculations. The linking of localizations between GT and S1/S2 is repeated, as localizations can be shifted in/out of the maximum linking distance due to the median offset. Of this linked dataset, the Jaccard index is calculated as follows:

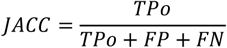

where *TPo, FP*, and *FN* are the true positive, false positive, and false negative localizations, respectively.

Then, only localizations that are present in all three datasets (GT, S1 and S2) are selected, and of these localizations, the root mean square error (RMSE) in a single dimension is calculated as follows:

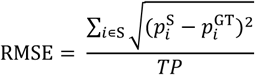

where *p*_*i*_ indicates the position of localization *i* in any dimension, and *S* indicates S1 or S2.

For saddle-point (SP), 1000 PSFs for every signal-to-noise combination were simulated (section 2.3), after which a calibration curve was created in SMALL-LABS-ct-pSMLM or SMAP with PSFs containing 3·10^4^ photons on a 1 photon/pixel background. For SMAP localizations, obtained localizations well outside the expected regime (10 pixels or further removed from center) were discarded, and frames containing multiple or no localizations were fully discarded. Note, localizations obtained with SMAP that were clearly misfitted (an offset in *z* by at least 3 times the average *z* offset calculated by ct-pSMLM) were discarded; no such discarding was performed for ct-pSMLM. For both ct-pSMLM and SMAP, the *x, y* and *z* positions were compared with the ground-truth, and the standard deviation of this offset was calculated for every intensity and background combination and is shown in the results. We note that the mean of the offset was centered around 0 for every tested intensity/noise/software combination.

### 2.6 Single-molecule microscopy

For SMLM experiments, we used a home-built super-resolution microscope similar to one reported previously [24]. Briefly, light from a fiber-coupled 642 nm laser (Omicron, Germany) was collimated using an achromatic lens (*f* = 30mm, Thorlabs) and conducted to a parabolic mirror (RC12APC-P01, Thorlabs). The laser light was then focused using an achromat lens (f = 150mm, Thorlabs) in front of a polychroic mirror (ZT532/640rpc, Chroma) into the backfocal plane of an 100x oil-immersion objective (CFI Plan Apo, NA = 1.45, Nikon Japan) such that a highly inclined illumination (HiLo) profile with a total laser power of ∼70 mW was achieved. Emitted fluorescence passing the objective, the polychroic mirror and a bandpass filter (ZET532/640m-TRF, Chroma) was then guided into a 4f geometry using the following lenses (1: *f* = 200mm, 2: *f* = 100mm, 3: *f* = 100mm) towards a Prime 95B sCMOS camera (Photometrics, Tucson, AZ, USA), resulting in an effective 115 by 115 nm pixel size. A deformable mirror (DMP40-P01, Thorlabs) was placed in the Fourier plane between lens 2 and 3. The deformable mirror used 40 segments with bending arms for tip-tilt control to modulate and introduce different Zernike modes. After calibrating (flattening) the deformable mirror via the REALM software (https://github.com/MSiemons/REALM and https://github.com/HohlbeinLab/Thorlabs_DM_Device_Adapter) we used the Zernike polynomials 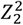 (primary astigmatism) and 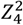 (secondary astigmatism) to induce saddle-point PSFs [38,40].

### 2.7 STORM experiment

A SAFe sample containing immobilized Cos7 fibroblasts from Green African Monkeys (ATCC) with Alexa Fluor 647 labeled tubulins was purchased from Abbelight (Paris, France). A nitrogen-flushed buffer containing 50 mM TRIS pH8, 10 mM NaCl, 10% glucose, 50 mM 2-mercaptoethanol, 68 µg/mL catalase, and 200 µg/mL glucose oxidase [27] was added to the sample chamber which was sealed off before the measurements. 60.000 frames of 20 ms length were recorded using the setup described in section 2.6. Analysis of the single-molecule data was performed as specified in section 2.4.

## 3. Methods

### 3.1 Principles of engineered PSF localization with pSMLM: Double-helix: DH-pSMLM

To localize double-helix (DH) PSFs, we rely on the fact that pSMLM-2D provides accurate lateral localization even when using a relatively small ROI around the center of an emitter [31]. Therefore, the two lobes rotating around each other (Fig. 1a) can be localized separately. During calibration, the distance and rotation between the two lobes is plotted against the axial position (Fig. 1b). The rotation is fitted with a third-order polynomial. This polynomial is weighted on the inverse of the standard deviation of each axial position if more than one calibration bead has been used.

**Fig. 1.**
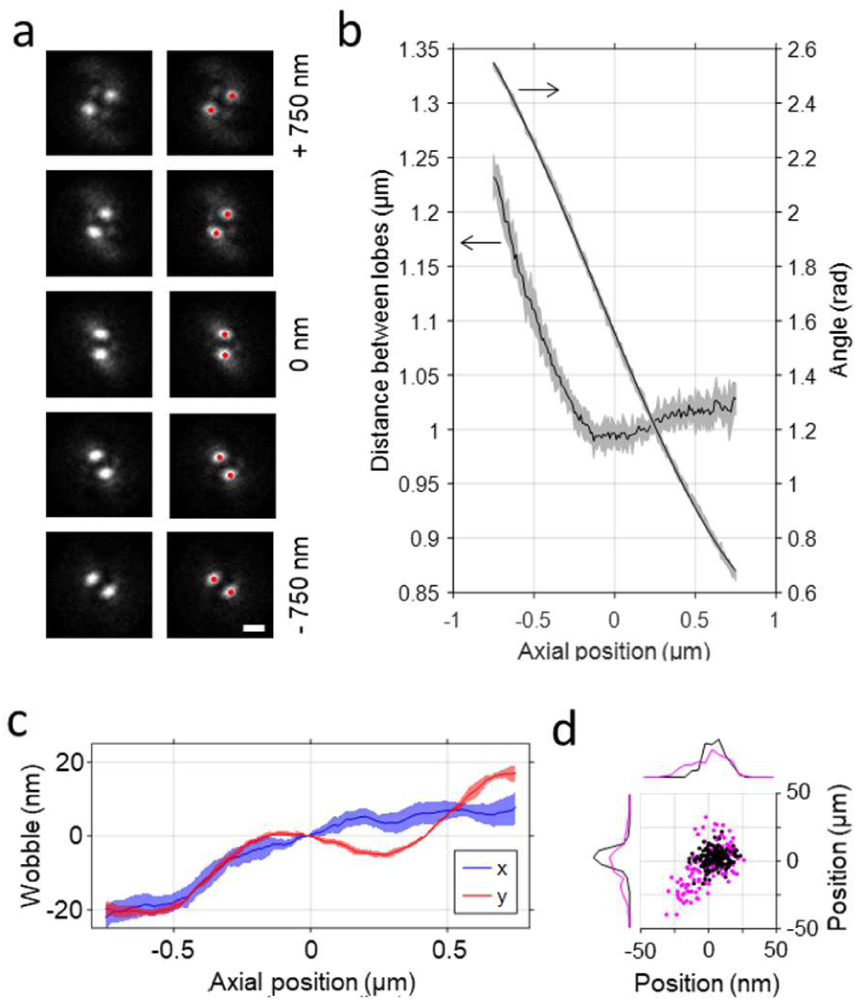
Double Helix PSF fitting via phasor-based localization. **a** Typical double helix axial position profile [41]. Red markers indicate initial 2D fits of single lobes by pSMLM. Scalebar represents 1 µm. **b** Typical calibration curve determined via DH-pSMLM. The distance between the lobes is used to link single lobes during analysis, while the angle between the lobes is used to calculate the axial position. **c** Typical wobble calculated from calibration. The solid line represents the average wobble in *x* or *y* at every axial position, the shaded area represents the standard deviation. **d** Wobble correction effect for a single calibration emitter. Magenta shows uncorrected lateral position of a fixed simulated emitter, while black shows the wobble-corrected position.

The lateral position is calculated as being the average lateral position of the two lobes, corrected for a ‘wobble’ factor. This wobble factor is determined in *x* and *y* as function of the emitters axial position (Fig. 1c) by comparing the lateral localization at all axial positions with the lateral localization at the axial center of the calibration dataset. The average of this wobble effect over an axial sliding window (user-defined, default value is set to 5 axial positions) is determined during calibration and stored for future correction of lateral localization calculation (Fig. 1d).

To extract positional information, first a standard pSMLM-2D fitting is performed [31]. The localizations in each frame are compared with each other to find pairs within the expected distance regime (determined during calibration; minimum and maximum of distance between lobe centers, with a ∼10% error margin), and are discarded if no pair can be found. During the linking of the lobes, priority is given to lobes that only have a single possible counter-lobe over those that have multiple options to reduce mis-fitting of closely positioned DH PSFs. The axial position is then determined from the rotation of the two lobes via the calibration curve. The obtained distance between lobes is checked against the distance determined during calibration at the found axial position, and the localization is discarded if these values differ more than ∼100 nm (user defined). Lastly, the lateral position is determined from the mean of the 2D-determined position of the two lobes, and corrected for the wobble determined during calibration (Fig 1d).

### 3.2 Principles of engineered PSF localization with pSMLM: Saddle-point and tetra-pod: ct-pSMLM

We analyze saddle-point (SP) and tetra-pod (TP) PSFs with an adapted phasor-based localization methodology. SP and TP have similar characteristics and show separation of a single point when in focus into two lobes above and below the focus in perpendicular directions [38,39]. Moreover, they are based on similar PSF deformations introduced by primary and secondary astigmatism Zernike coefficients [40].

We modified a spectral phasor-based approach [48] in which the convolution of arbitrary profiles in real space is a linear combination of their respective phasor representations in phasor space. In this approach, the normalized intensity ratio between the original profiles in the convoluted profile (real space) is represented as the distance of the original phasor profiles to the convoluted phasor profile (phasor space). This entails that if two profiles are combined with a 1:1 ratio, the convoluted phasor representation is on the mid-point of the line between the phasor representations of the original profiles.

In SP and TP PSFs, the final spatial representation of the PSF is a convolution of two separated lobes of identical intensity. Thus, SP and TP PSFs can be treated as a 1:1 ratio of arbitrary profiles that are separated at a varying distance, depending on the axial position of the emitters. Note that by orientating the respective optical components correctly, this separation can be achieved perfectly on the *x*- or *y*-axis. Therefore, the value for the separation *d*_lobes_, along with the orientation of this separation provides suitable information for calibration of SP and TP PSFs. To determine the separation *d*_lobes_ with phasor-based single-molecule localization microscopy (pSMLM), we assume that the width of the individual lobes in the direction of the convolution is identical to the width of the convoluted PSF in the other, unconvoluted spatial direction. For illustration, we show a combination of two 2-dimensional Gaussian distributions (Fig. 2a,b,c). The phasor representation of the individual Gaussian distributions is represented by a single phasor for both dimensions, each having a certain, but different, angle representing the emitter’s position in real space [31].

**Fig. 2.**
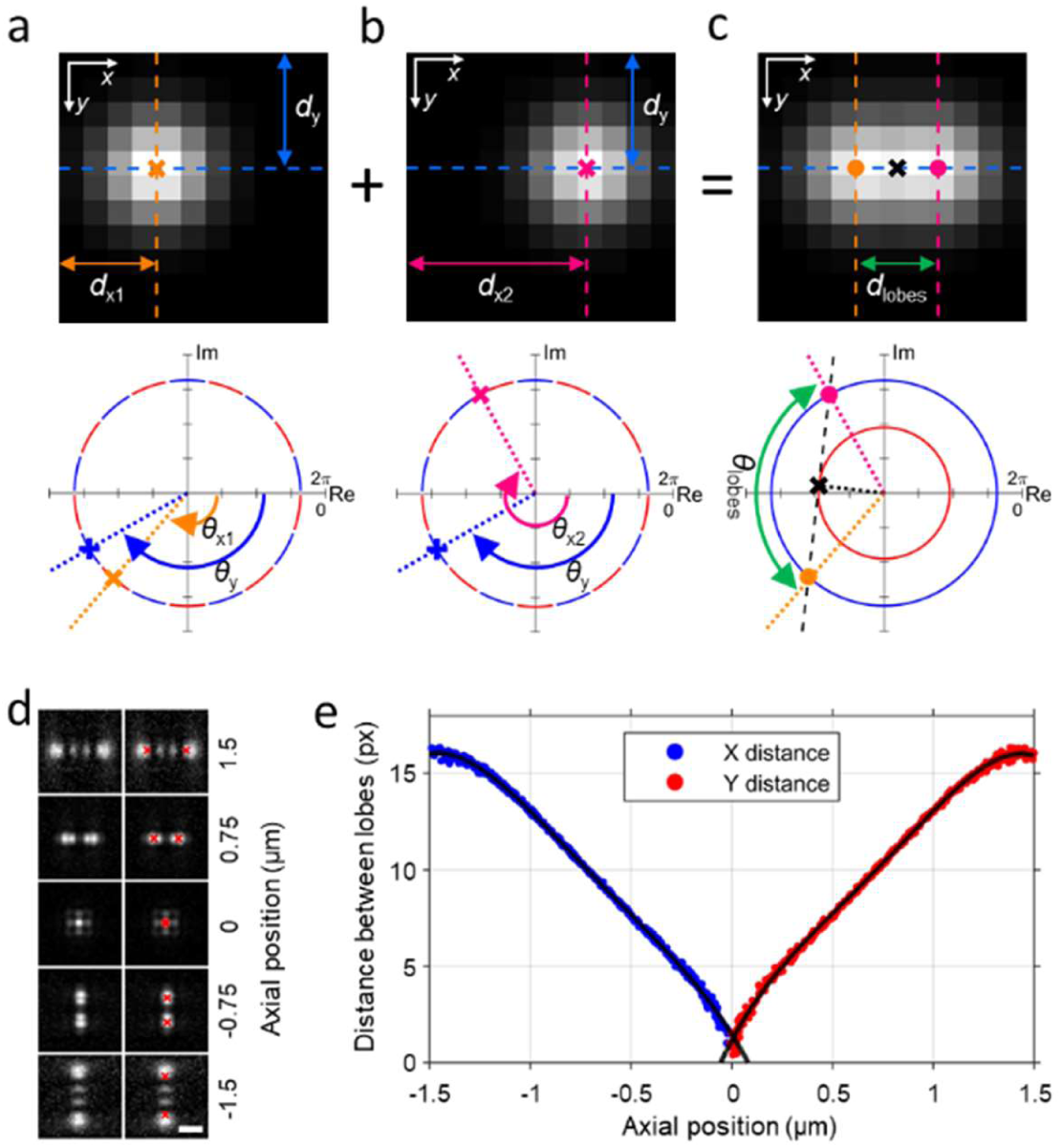
Principle of circular-tangent phasor-based single-molecule localization microscopy (ct-pSMLM) which can be used for saddle-point (SP) and tetra-pod (TP) PSF localization. **a** Single simulated 2D-Gaussian profile (top). The phasor representation (bottom) in the *x* dimension shows a single phasor, of which the angle *θ*x1 represents *x*-position *d*x1 of the emitter in real space (orange cross in both real space and phasor space). The angle *θ*y in phasor space (blue plus) represents *y*-position *d*y in real space. The magnitude of the *x*-dimension (red) and *y-*dimension (blue) are shown as circles with equal radius. **b** Identical to **a**, but with the 2D-Gaussian profile at a different position. **c** Top: Combination of the two Gaussian profiles shown in **a** and **b**. The wider *x* profile corresponds to a smaller *x* phasor magnitude (red circle) as compared to the profile and magnitude in *y*. Bottom: The phasor representation of the profile in *x* is represented by the black cross (the *y*-phasor is omitted for clarity). Next, a line (black dashed) is placed perpendicular on the *x* phasor magnitude circle. The positions where this line crosses the *y* phasor magnitude circle are indicated by the orange and magenta dots in phasor space. These values are normalized values for the single emitter positions in real space, indicated by orange and magenta dots. The angle between the two phasor angles is *θ*lobes (green) and represents *d*lobes in real space. **d** Representative simulated SP emitters at varying axial positions (see Methods). Red markers indicate the obtained lobe positions via ct-pSMLM. Scalebar represents 1 µm. **e** Typical calibration curve in which the separation of the lobes in *x* and *y* is plotted as a function of the *z* position.

Then, if reasoned from the convoluted PSF (Fig. 2c) to obtain the individual lobes, the tangent at the magnitude circle in the convoluted spatial dimension (broad spatial dimension; small phasor magnitude; black cross located on the red circle in Fig. 2c) will intersect the magnitude of the smaller spatial dimension (large phasor magnitude; represented as a blue circle) at two points (Fig. 1c; magenta and orange dots). These points are a measure for original arbitrary profiles with identical spatial sizes in both dimensions that combine in a 1:1 intensity ratio to result in the convoluted profile. The angle between these two obtained intersectional points (*θ*_lobes_) in phasor space is a direct normalized value for the distance *d*_lobes_ in real space (Fig. 2c). We call this method circular-tangent pSMLM (ct-pSMLM).

The obtained *d*_lobes_ is used to create well-defined calibration curves for both SP and TP (SP shown in Fig. 2d,) that can be fitted with arbitrary functions (e.g. a fourth-order polynomial) to deduce axial positional information from experimental PSFs (Fig. 2e). The lateral localization information of the SP or TP PSFs is still inherently present in the original phasor-representation of the complete PSF.

Determining localization of the SP or TP PSFs in the SMALL-LABS-pSMLM software consists of two parts: finding the central positions and further analysis with ct-pSMLM. The mid-points of single PSFs are determined by first checking whether two detected emitters that could represent two lobes of a single SP/TP PSF belong to the same PSF. If these emitters have little deviation in one dimension (<0.5 px) and are slightly separated in the other dimension (less than the calibrated maximum distance), the mid-point of these emitters is calculated and stored. If no other lobe can be found, it is assumed the located emitter is the mid-point of the SP/TP PSF. Then, ct-pSMLM is performed around the central point with a reasonably large region of interest (> 2 µm) to obtain *d*_lobes_ and to calculate the axial position.

## 4. Results

### 4.1 Double-helix

To evaluate the performance of DH⍰pSMLM, we performed fitting of simulated datasets [44] via the full pSMLM-updated SMALL-LABS software package and compared with the currently best performing non-machine learned localization algorithm (experimental PSF spline fitting methodology incorporated in SMAP [26]).

As the ground truth of these datasets is publicly available, we were able to extract (Table 1) quantitative performance parameters such as the expected deviation of localization accuracy in all three dimensions (root mean squared error, RMSE) and the Jaccard index JACC (a measure for correctly and incorrectly localized particles [41]). These performance parameters are calculated from localizations that were found in both software packages and in the Ground-Truth datasets.

**Table 1.**
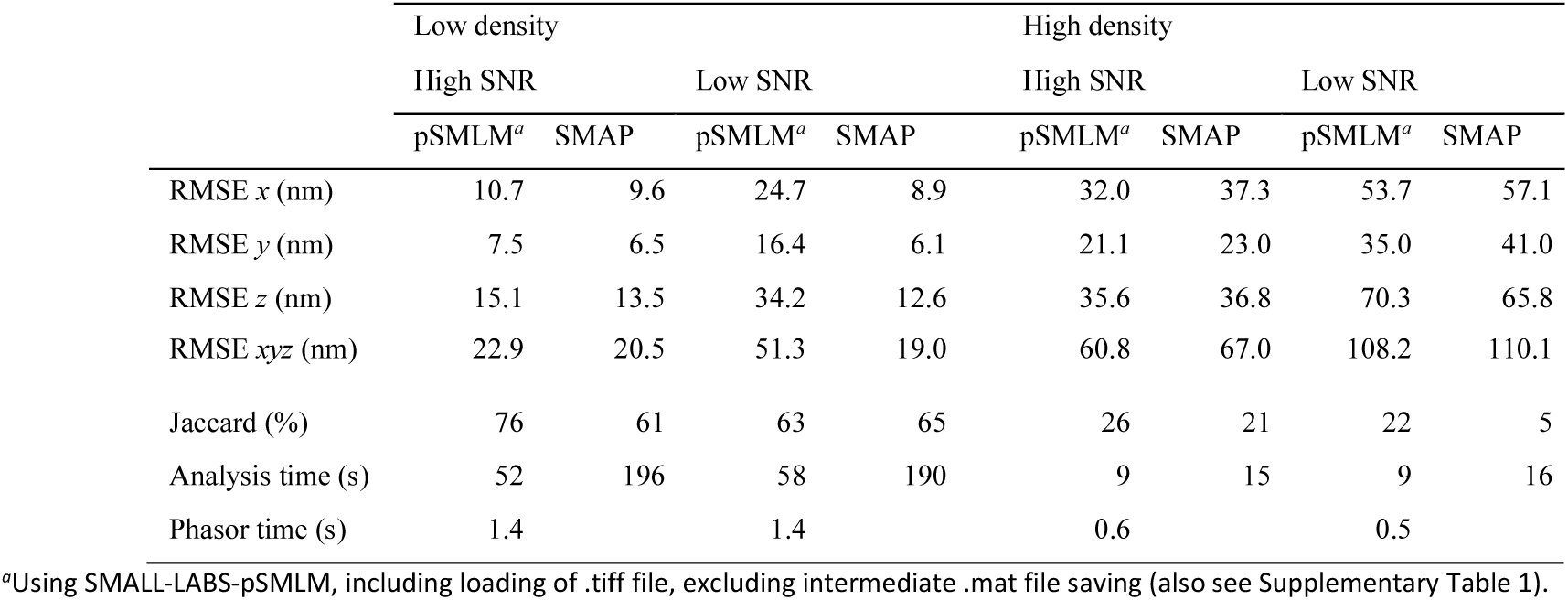
Double Helix PSF fitting performance, comparing SMALL-LABS-pSMLM with SMAP.

|  | Low density |  |  |  | High density |  |  |  |
| --- | --- | --- | --- | --- | --- | --- | --- | --- |
|  | High SNR |  | Low SNR |  | High SNR |  | Low SNR |  |
|  | pSMLM<sup>a</sup> | SMAP | pSMLM<sup>a</sup> | SMAP | pSMLM<sup>a</sup> | SMAP | pSMLM<sup>a</sup> | SMAP |
| RMSE $x$ (nm) | 10.7 | 9.6 | 24.7 | 8.9 | 32.0 | 37.3 | 53.7 | 57.1 |
| RMSE $y$ (nm) | 7.5 | 6.5 | 16.4 | 6.1 | 21.1 | 23.0 | 35.0 | 41.0 |
| RMSE $z$ (nm) | 15.1 | 13.5 | 34.2 | 12.6 | 35.6 | 36.8 | 70.3 | 65.8 |
| RMSE $xyz$ (nm) | 22.9 | 20.5 | 51.3 | 19.0 | 60.8 | 67.0 | 108.2 | 110.1 |
| Jaccard (%) | 76 | 61 | 63 | 65 | 26 | 21 | 22 | 5 |
| Analysis time (s) | 52 | 196 | 58 | 190 | 9 | 15 | 9 | 16 |
| Phasor time (s) | 1.4 |  | 1.4 |  | 0.6 |  | 0.5 |  |

<sup>a</sup>Using SMALL-LABS-pSMLM, including loading of .tiff file, excluding intermediate .mat file saving (also see Supplementary Table 1).

We observe that both SMALL-LABS-pSMLM and SMAP have comparable RMSE errors (Table 1) in the order of 10-25 nm for the low density (LD), high signal to noise (SNR) dataset which are similar to the ones reported previously for SMAP [26,41]. However, at low signal to noise levels, SMAP outperforms SMALL-LABS-pSMLM on all performance indicators. This is presumably due to SMAP using the full PSF at once, while SMALL-LABS-pSMLM splits localization in two steps. This results in SMALL-LABS-pSMLM working with a lower apparent signal to noise level, causing a lower localization accuracy. We note that the reported RMSE values for SMAP analysis of the LD, low SNR dataset are counter-intuitively better than those of SMAP analysis of the LD, high SNR dataset. This is a result of the RMSE calculation methodology used (Material and methods), as only localizations that are found in both software analyses as well as in the ground-truth are used for RMSE calculations.

We observe that SMALL-LABS-pSMLM outperforms SMAP in terms of localization recall rates (Jaccard index, Table 1) at high SNR (23% increased), but not at low SNR (4% decrease). The Jaccard values for SMAP are slightly lower than reported earlier [41] (Material and methods), but can be compared directly with the Jaccard values for SMALL-LABS-pSMLM reported here. We note that no background subtraction is performed in SMAP, while SMALL-LABS-pSMLM subtracts the background based on foreground temporal variations (SMALL-LABS [43]).

Both software packages are not capable of recognizing HD PSFs with a good recall rate, although SMALL-LABS-pSMLM outperforms SMAP in all conditions, as single DH lobes are localized with only 5 × 5 pixel ROIs, decreasing the influence of the other nearby emitters.

SMALL-LABS-pSMLM is ∼3 – 4 x faster compared to SMAP for low density datasets, and ∼2x faster for high density datasets. However, most analysis time for SMALL-LABS-pSMLM (∼65%) is spend on the background correction and format conversion rather than approximate localization or sub-pixel DH⍰pSMLM localization (Supplementary Table 1). The localization procedure itself can achieve 1 - 1.5·10^4^ localizations per second on a standard CPU.

### 4.2 Saddle-point and tetra-pod

The performance of the localization of saddle-point (SP) PSFs was assessed and compared to experimental PSF spline fitting [26]. As ct-pSMLM is a non-iterative method, high localization rates of up to 2.5·10^5^ localizations per second were achieved on standard CPUs (Fig. 3a). This is an order of magnitude lower than traditional pSMLM-3D [31], mostly due to the large required region of interest around the PSF (>2 µm; here 23 × 23 px), and partly due to the additional computations required for ct-pSMLM. Taken alone, the additional computations of ct-pSMLM compared to pSMLM-3D only result in a 10 - 40% decrease in localization rates (∼10% for at a large region of interests 23 × 23 px; ∼40% decrease for 7 × 7 px).

**Fig. 3.**
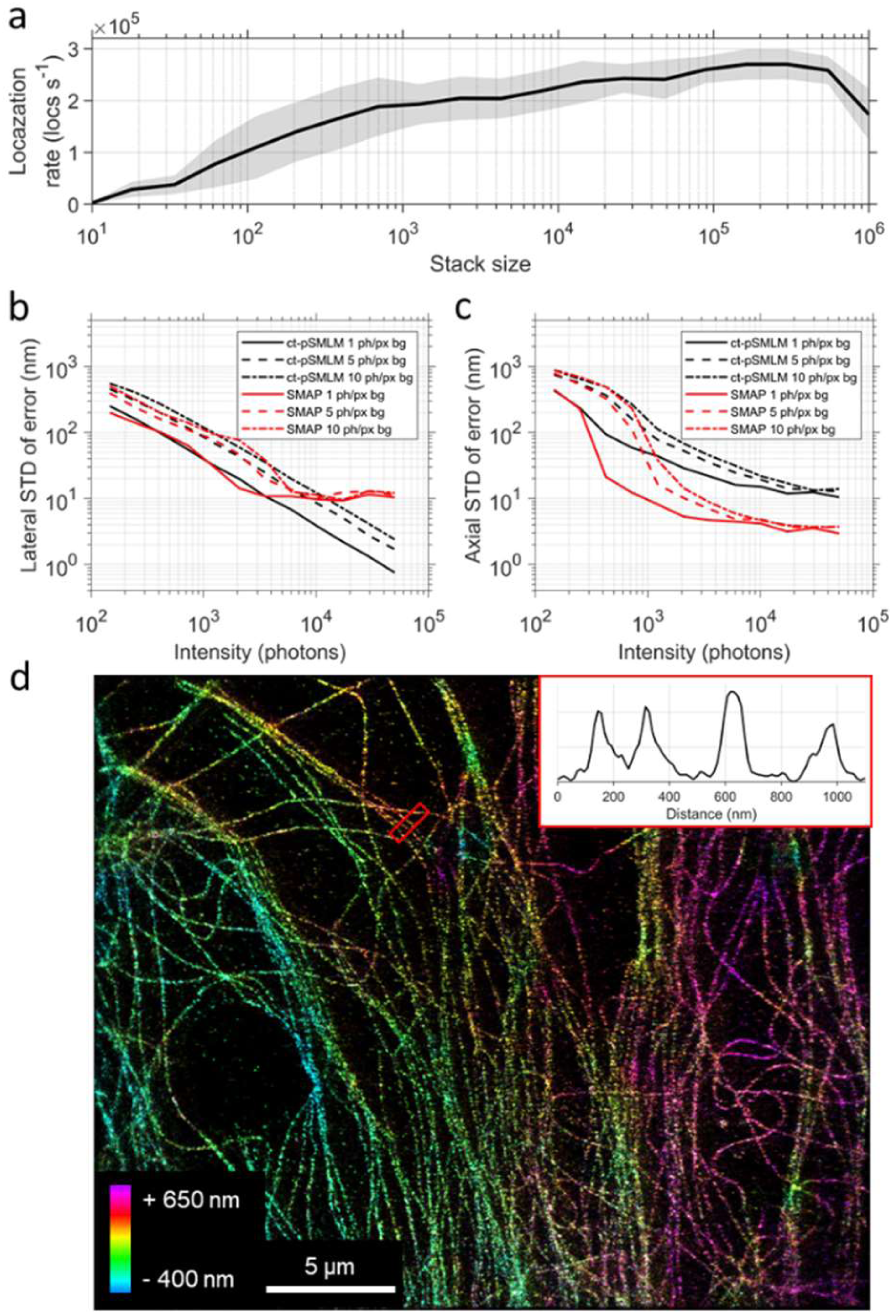
Saddle-point (SP) ct-pSMLM performance. **a** Speed of ct-pSMLM, based on a 23 × 23 pixel image stack. Shaded area indicates the standard deviation. **b, c** Localization precision of ct-pSMLM, based on simulated saddle-point point spread functions with different intensities (*x* axis) and background noise levels (line types). Lateral localization precision is shown in b, while axial precision is shown in c. **d** Experimental STORM microtubule network analyzed with ct-pSMLM integrated in SMALL-LABS-pSMLM. Inset: Lateral profile of the red boxed outline.

The lateral localization accuracy of ct-pSMLM is in line with that of experimental PSF spline fitting (Fig 3b), and decreases from ∼100 nm (∼1 pixel) at typical (∼200 - 1300) photon values for fluorescent proteins to ∼10 nm (∼0.1 pixels) at typical (∼(2 – 11) × 10^3^) photon values for organic fluorophores. The localisation accuracy is roughly one order of magnitude lower than the lateral localization accuracy of non-engineered PSFs at high photon values (∼0.08 pixels and ∼0.01 pixels, respectively [31]), and roughly 1.5x worse than AS PSFs (∼0.05 pixels [31]) caused by lower effective signal to noise ratio due to the expanded PSF. We observed a lower limit in lateral localization accuracy for SMAP fitting of ∼10 nm (∼0.1 pixel), which has an unknown origin.

The axial localization accuracy of ct-pSMLM increases with increasing photon values as well (Fig 3c). The average axial accuracy at typical photon values for organic fluorophores is ∼40 nm, which is ∼3x worse than AS PSFs with similar total photon counts and photons/pixel background [31]. The best obtainable axial accuracy is limited by the sub-optimal fitting of the calibration curve to around 11 nm, which is similar to AS PSFs (Fig 3c, SI fig 2, [31]). We attribute this lower axial accuracy of SP PSFs compared to AS PSFs again to lower effective signal to noise ratios due to expanded PSFs. Up to 20% of SMAP-localized emitters had to be discarded from calculating the *z* offset, as these were substantially misfitted (> 3 times the corresponding ct-pSMLM *z* offset, see Methods). We furthermore demonstrate the implementation of ct-pSMLM in SMALL-LABS-pSMLM by analysing an experimental STORM experiment showing labelled microtubule of a Monkey cell line (Fig 3d, Material and methods). The total analysis time for the SMALL-LABS-pSMLM analysis for the ∼8 GB, 60.000 frames dataset containing 1.5 million localizations was ∼15 minutes on a standard CPU, including file conversion (Supplementary Table 1), of which the ct-pSMLM sub-pixel fitting routine comprised just 77 seconds.

## 5. Discussion

Here we present additions on the phasor-based single-molecule localization microscopy (pSMLM) framework to localize double helix (DH), saddle-point (SP), and tetra-pod (TP) PSFs with very good accuracy and speed on standard CPUs. In the current implementation, DH-pSMLM can achieve up to 1.5·10^4^ localizations/second on a 3.10 GHz processing unit, while ct-pSMLM, the basis for SP and TP localization, can achieve up to 2.5·10^5^ localizations/second. Specifically, ct-pSMLM is designated for real-time localization methods, combined with computationally inexpensive filtering and background subtraction methods, to better enable (automated) feedback-oriented SMLM instrumentation. Possibly a pSMLM-based methodology could be implemented on the integrated circuits of cameras to further increase end-user accessibility of advanced single-molecule techniques.

The DH-pSMLM implementation in the SMALL-LABS software has similar performance as the current state-of-the-art methods when using organic fluorophores, while decreasing the overall analysis time when ran on a CPU. We note that our implementation is not particularly sensitive to overfitting, as stringent constraints on the pair-finding are used. This allows for some false positives during the initial localization steps which are later discarded.

Ct-pSMLM improves on our previous phasor implementation [31] by offering a direct way of determining the distance between two emission peaks of a single PSF, well-suited for the quantification of the axial position in SP and TP PSFs. Here, perfect horizontal and vertical elongation of the PSFs is a requirement for ct-pSMLM to perform. Our algorithm is capable of retrieving the emitters location with a precision similar to current best non-machine learning localization algorithm [41], and is mostly limited by the fitting of the calibration curve. Naturally, for ct-pSMLM to work correctly, no other emitters or highly inhomogeneous background should be present in the fitting region. As SP and TP PSFs require large ROIs (∼23 × 23px), this results in substantially lower accessible emitter density compared to approaches using standard and astigmatic PSFs. For high-density engineered PSF localization approaches, we point to alternative approaches such as deep learning [49,50] or matching pursuit [51].

We incorporated the novel pSMLM-derivative localization methodologies in the SMALL-LABS software [43]. The updated SMALL-LABS-pSMLM software package expands the original work with a user-friendly GUI, wavelet filtering, drift-correction in 3D, and result image generation. We believe that the software package strikes an excellent balance between fast analysis, accurate results, experimental freedom, good expandability, and hassle-free installation and operation. The software is freely available at https://github.com/HohlbeinLab/SMALL-LABS-pSMLM.

## Funding

K.J.A.M was funded by a VLAG Ph.D. fellowship awarded to J.H. This work is part of the research programmes LICENSE and LocalBioFood with project numbers 731.017.301 and 731.017.204, which are financed by the Dutch Research Council (NWO).

## Acknowledgements

We gratefully acknowledge Bernd Rieger and Sjoerd Stallinga (TU Delft) for sharing MATLAB code to simulate engineered point spread functions. We thank Julie Biteen, Laurent Geffroy, and Jonas Ries for helpful discussions on software. We thank Arjen Bader for fruitful discussions on applications of phasor analysis on single molecule localization microscopy.

## Author contributions

Conceptualization: KM and JH. Data curation: KM, AJ, SY. Formal analysis: KM. Funding acquisition: JH. Investigation: KM, AJ, SY. Methodology: KM and JH. Project administration: JH. Software: KM. Supervision: JH. Visualization: KM. Writing – original draft: KM. Writing – review & editing: All authors.

## Supplementary information

**Supplementary Table 1.** Detailed analysis time for double-helix fitting by pSMLM-SL-DH

|  | TIFF<br>conversion | Intermediate<br>file saving | Background<br>subtraction | Total analysis<br>time (s) | pSMLM-DH<br>fitting time (s) |
| --- | --- | --- | --- | --- | --- |
| N1-LD<br>(20,000 frames) | Yes | Yes | Yes | $72.7 \pm 1.7$ | 1.4 |
| | Yes | Yes | No | $43.7 \pm 1$ | 1.4 |
| | Yes | No | Yes | $51.8 \pm 0.4$ | 1.4 |
| | Yes | No | No | $34.6 \pm 0.8$ | 1.4 |
| | No | Yes | Yes | $57 \pm 1.1$ | 1.4 |
| | No | Yes | No | $25.8 \pm 0.3$ | 1.4 |
| | No | No | Yes | $42 \pm 0.2$ | 1.4 |
| | No | No | No | $24.6 \pm 0.2$ | 1.4 |
| N1-HD<br>(2,000 frames) | Yes | Yes | Yes | $13.4 \pm 0.3$ | 0.5 |
| | Yes | Yes | No | $5.8 \pm 0.3$ | 0.6 |
| | Yes | No | Yes | $8.5 \pm 0.0$ | 0.6 |
| | Yes | No | No | $4.2 \pm 0.2$ | 0.6 |
| | No | Yes | Yes | $11.6 \pm 0.1$ | 0.6 |
| | No | Yes | No | $3.9 \pm 0.0$ | 0.6 |
| | No | No | Yes | $8.2 \pm 0.0$ | 0.6 |
| | No | No | No | $3.8 \pm 0.1$ | 0.6 |

**S. Fig. 1.**
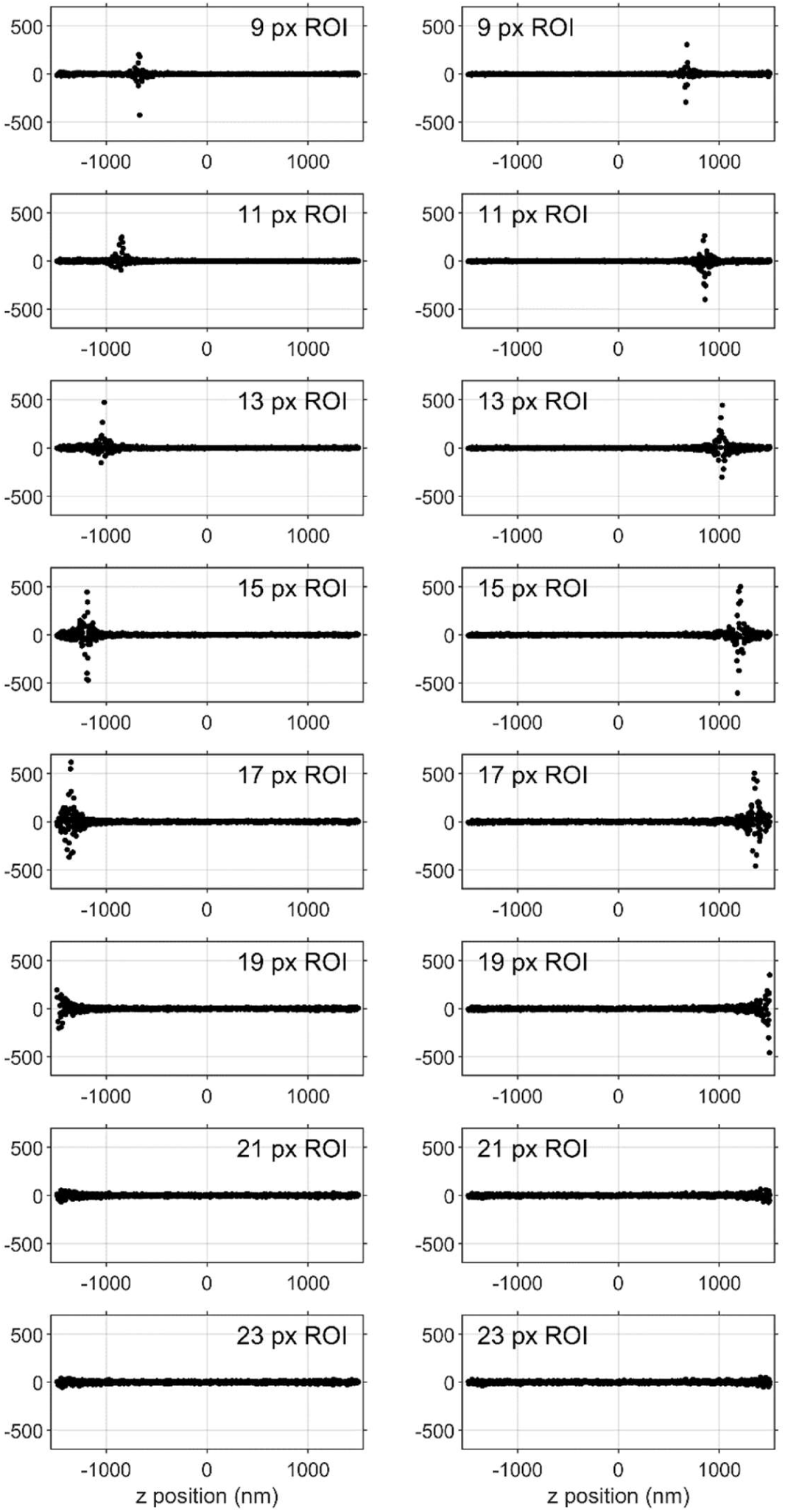
Offset (in nm) in *y* (left) and *x* (right) based on *z* position (x-axis) and ROI (vertical panels) used for ct-pSMLM. Note that the offset error is in the direction of the lobes (*y*-direction at negative *z* positions, *x*-direction at positive *z* positions), is very localized, and shifts with increasing ROI size towards more extreme *z* positions.

**S. Fig. 2.**
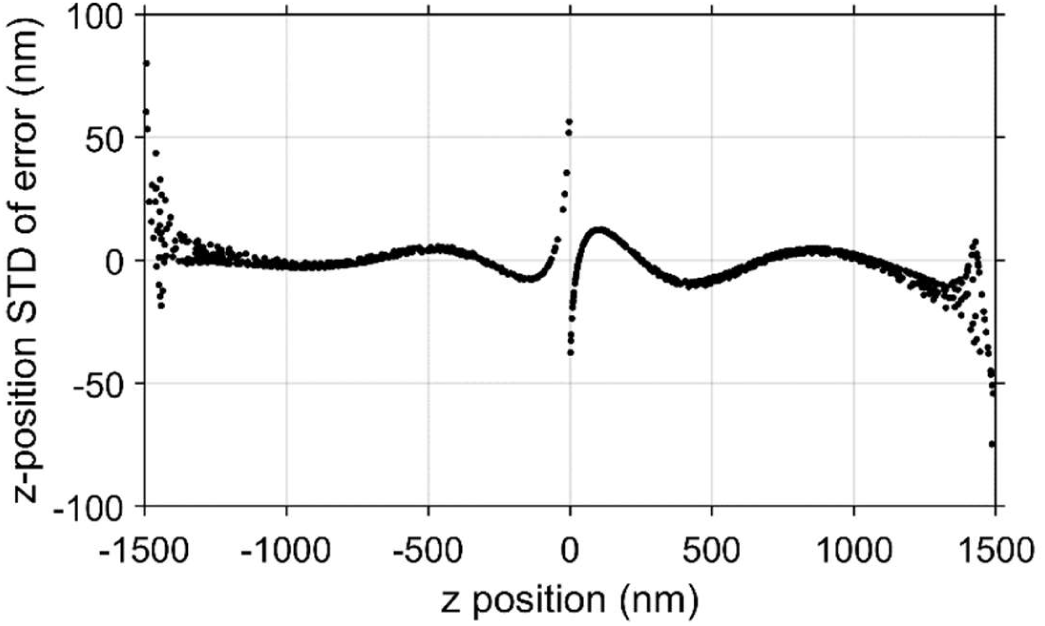
Lateral accuracy of ct-pSMLM as a function of z position, calculated via simulated PSFs without introduced noise. The mismatch between the fitted calibration curve and the actual distance is an effective lower limit of the lateral localization accuracy, and is on average 11 nm for our tested simulated PSFs.

## Notes

### Competing Interest Statement

The authors have declared no competing interest.

## References

1. L. Schermelleh, A. Ferrand, T. Huser, C. Eggeling, M. Sauer, O. Biehlmaier, and G. P. C. Drummen, “Super-resolution microscopy demystified,” Nature Cell Biology 21(1), 72 (2019).

2. J. Vangindertael, R. Camacho, W. Sempels, H. Mizuno, P. Dedecker, and K. P. F. Janssen, “An introduction to optical super-resolution microscopy for the adventurous biologist,” Methods and applications in fluorescence 6(2), 022003 (2018).

3. M. Heilemann, “Fluorescence microscopy beyond the diffraction limit,” Journal of Biotechnology 149(4), 243–251 (2010).

4. M. J. Rust, M. Bates, and X. Zhuang, “Sub-diffraction-limit imaging by stochastic optical reconstruction microscopy (STORM),” Nature methods 3(10), 793 (2006).

5. M. Heilemann, S. van de Linde, M. Schüttpelz, R. Kasper, B. Seefeldt, A. Mukherjee, P. Tinnefeld, and M. Sauer, “Subdiffraction-Resolution Fluorescence Imaging with Conventional Fluorescent Probes,” Angewandte Chemie International Edition 47(33), 6172–6176 (2008).

6. E. Betzig, G. H. Patterson, R. Sougrat, O. W. Lindwasser, S. Olenych, J. S. Bonifacino, M. W. Davidson, J. Lippincott-Schwartz, and H. F. Hess, “Imaging intracellular fluorescent proteins at nanometer resolution,” Science 313(5793), 1642–1645 (2006).

7. M. G. Gustafsson, “Surpassing the lateral resolution limit by a factor of two using structured illumination microscopy,” Journal of microscopy 198(2), 82–87 (2000).

8. S. W. Hell and J. Wichmann, “Breaking the diffraction resolution limit by stimulated emission: stimulated-emission-depletion fluorescence microscopy,” Optics letters 19(11), 780–782 (1994).

9. S. W. Hell, “Toward fluorescence nanoscopy,” Nature biotechnology 21(11), 1347 (2003).

10. T. Dertinger, R. Colyer, G. Iyer, S. Weiss, J. Enderlein, and J. W. Sedat, “Fast, Background-Free, 3D Super-Resolution Optical Fluctuation Imaging (SOFI),” Proceedings of the National Academy of Sciences of the United States of America 106(52), 22287–22292 (2009).

11. N. Gustafsson, S. Culley, G. Ashdown, D. M. Owen, P. M. Pereira, and R. Henriques, “Fast live-cell conventional fluorophore nanoscopy with ImageJ through super-resolution radial fluctuations,” Nature Communications 7, 12471 (2016).

12. F. Balzarotti, Y. Eilers, K. C. Gwosch, A. H. Gynnå, V. Westphal, F. D. Stefani, J. Elf, and S. W. Hell, “Nanometer resolution imaging and tracking of fluorescent molecules with minimal photon fluxes,” Science 355(6325), 606–612 (2017).

13. C. S. Smith, N. Joseph, B. Rieger, and K. A. Lidke, “Fast, single-molecule localization that achieves theoretically minimum uncertainty,” Nat Meth 7(5), 373–375 (2010).

14. E. Abbe, “Beiträge zur Theorie des Mikroskops und der mikroskopischen Wahrnehmung,” Archiv für mikroskopische Anatomie 9(1), 413–418 (1873).

15. A. von Diezmann, Y. Shechtman, and W. E. Moerner, “Three-Dimensional Localization of Single Molecules for Super-Resolution Imaging and Single-Particle Tracking,” Chem. Rev. 117(11), 7244–7275 (2017).

16. S. Stallinga and B. Rieger, “Accuracy of the Gaussian point spread function model in 2D localization microscopy,” Optics express 18(24), 24461–24476 (2010).

17. M. Sauer and M. Heilemann, “Single-Molecule Localization Microscopy in Eukaryotes,” Chem. Rev. 117(11), 7478–7509 (2017).

18. D. Baddeley and J. Bewersdorf, “Biological Insight from Super-Resolution Microscopy: What We Can Learn from Localization-Based Images,” Annu. Rev. Biochem. 87(1), 965–989 (2018).

19. U. Endesfelder, “From single bacterial cell imaging towards in vivo singlemolecule biochemistry studies,” Essays In Biochemistry EBC20190002 (2019).

20. M. Stracy, C. Lesterlin, F. G. de Leon, S. Uphoff, P. Zawadzki, and A. N. Kapanidis, “Live-cell superresolution microscopy reveals the organization of RNA polymerase in the bacterial nucleoid,” PNAS 112(32), E4390–E4399 (2015).

21. M. Stracy and A. N. Kapanidis, “Single-molecule and super-resolution imaging of transcription in living bacteria,” Methods 120, 103–114 (2017).

22. J. N. A. Vink, K. J. A. Martens, M. Vlot, R. E. McKenzie, C. Almendros, B. E. Bonilla, D. J. W. Brocken, J. Hohlbein, and S. J. J. Brouns, “Direct visualization of native CRISPR target search in live bacteria reveals Cascade DNA surveillance mechanism,” bioRxiv 589119 (2019).

23. B. Turkowyd, H. Müller-Esparza, V. Climenti, N. Steube, U. Endesfelder, and L. Randau, “Live-cell single-particle tracking photoactivated localization microscopy of Cascade-mediated DNA surveillance.,” Methods in enzymology 616, 133–171 (2019).

24. K. J. A. Martens, S. P. B. van Beljouw, S. van der Els, J. N. A. Vink, S. Baas, G. A. Vogelaar, S. J. J. Brouns, P. van Baarlen, M. Kleerebezem, and J. Hohlbein, “Visualisation of dCas9 target search in vivo using an open-microscopy framework,” Nat Commun 10(1), 3552 (2019).

25. J. V. Thevathasan, M. Kahnwald, K. Cieslinski, P. Hoess, S. K. Peneti, M. Reitberger, D. Heid, K. C. Kasuba, S. J. Hoerner, and Y. Li, “Nuclear pores as versatile reference standards for quantitative superresolution microscopy,” Nature Methods 16(10), 1045–1053 (2019).

26. Y. Li, M. Mund, P. Hoess, J. Deschamps, U. Matti, B. Nijmeijer, V. J. Sabinina, J. Ellenberg, I. Schoen, and J. Ries, “Real-time 3D single-molecule localization using experimental point spread functions,” Nature Methods 15(5), 367–369 (2018).

27. A. Jimenez, K. Friedl, and C. Leterrier, “About samples, giving examples: Optimized Single Molecule Localization Microscopy,” Methods (2019).

28. K. I. Mortensen, L. S. Churchman, J. A. Spudich, and H. Flyvbjerg, “Optimized localization analysis for single-molecule tracking and super-resolution microscopy,” Nature methods 7(5), 377 (2010).

29. R. Parthasarathy, “Rapid, accurate particle tracking by calculation of radial symmetry centers,” Nat Meth 9(7), 724–726 (2012).

30. H. P. Babcock and X. Zhuang, “Analyzing Single Molecule Localization Microscopy Data Using Cubic Splines,” Scientific Reports 7(1), 552 (2017).

31. K. J. A. Martens, A. N. Bader, S. Baas, B. Rieger, and J. Hohlbein, “Phasor based single-molecule localization microscopy in 3D (pSMLM-3D): An algorithm for MHz localization rates using standard CPUs,” The Journal of Chemical Physics 148(12), 123311 (2018).

32. C. Franke, M. Sauer, and S. van de Linde, “Photometry unlocks 3D information from 2D localization microscopy data,” nature methods 14(1), 41 (2017).

33. Y. Zhou, M. Handley, G. Carles, and A. R. Harvey, “Advances in 3D single particle localization microscopy,” APL Photonics 4(6), 060901 (2019).

34. B. Huang, W. Wang, M. Bates, and X. Zhuang, “Three-Dimensional Super-Resolution Imaging by Stochastic Optical Reconstruction Microscopy,” Science 319(5864), 810–813 (2008).

35. L. Holtzer, T. Meckel, and T. Schmidt, “Nanometric three-dimensional tracking of individual quantum dots in cells,” Appl. Phys. Lett. 90(5), 053902 (2007).

36. R. Henriques, M. Lelek, E. F. Fornasiero, F. Valtorta, C. Zimmer, and M. M. Mhlanga, “QuickPALM: 3D real-time photoactivation nanoscopy image processing in ImageJ,” Nat Meth 7(5), 339–340 (2010).

37. S. R. P. Pavani, M. A. Thompson, J. S. Biteen, S. J. Lord, N. Liu, R. J. Twieg, R. Piestun, and W. E. Moerner, “Three-dimensional, single-molecule fluorescence imaging beyond the diffraction limit by using a double-helix point spread function,” PNAS 106(9), 2995–2999 (2009).

38. Y. Shechtman, S. J. Sahl, A. S. Backer, and W. E. Moerner, “Optimal Point Spread Function Design for 3D Imaging,” Phys. Rev. Lett. 113(13), 133902 (2014).

39. Y. Shechtman, L. E. Weiss, A. S. Backer, S. J. Sahl, and W. E. Moerner, “Precise three-dimensional scan-free multiple-particle tracking over large axial ranges with tetrapod point spread functions,” Nano letters 15(6), 4194–4199 (2015).

40. A. Aristov, B. Lelandais, E. Rensen, and C. Zimmer, “ZOLA-3D allows flexible 3D localization microscopy over an adjustable axial range,” Nature Communications 9(1), 2409 (2018).

41. D. Sage, T.-A. Pham, H. Babcock, T. Lukes, T. Pengo, J. Chao, R. Velmurugan, A. Herbert, A. Agrawal, S. Colabrese, A. Wheeler, A. Archetti, B. Rieger, R. Ober, G. M. Hagen, J.-B. Sibarita, J. Ries, R. Henriques, M. Unser, and S. Holden, “Super-resolution fight club: assessment of 2D and 3D singlemolecule localization microscopy software,” Nature Methods 16(5), 387 (2019).

42. P. N. Petrov, Y. Shechtman, and W. E. Moerner, “Measurement-based estimation of global pupil functions in 3D localization microscopy,” Opt. Express, OE 25(7), 7945–7959 (2017).

43. B. P. Isaacoff, Y. Li, S. A. Lee, and J. S. Biteen, “SMALL-LABS: Measuring Single-Molecule Intensity and Position in Obscuring Backgrounds,” Biophysical Journal 116(6), 975–982 (2019).

44. I. Izeddin, J. Boulanger, V. Racine, C. G. Specht, A. Kechkar, D. Nair, A. Triller, D. Choquet, M. Dahan, and J. B. Sibarita, “Wavelet analysis for single molecule localization microscopy,” Opt. Express, OE 20(3), 2081–2095 (2012).

45. Y. Wang, J. Schnitzbauer, Z. Hu, X. Li, Y. Cheng, Z.-L. Huang, and B. Huang, “Localization events-based sample drift correction for localization microscopy with redundant cross-correlation algorithm,” Opt. Express, OE 22(13), 15982–15991 (2014).

46. M. Ovesný, P. Krížek, J. Borkovec, Z. Švindrych, and G. M. Hagen, “ThunderSTORM: a comprehensive ImageJ plug-in for PALM and STORM data analysis and super-resolution imaging,” Bioinformatics 30(16), 2389–2390 (2014).

47. D. W. Scott, “Averaged shifted histograms: effective nonparametric density estimators in several dimensions,” The Annals of Statistics 1024–1040 (1985).

48. F. Fereidouni, A. N. Bader, and H. C. Gerritsen, “Spectral phasor analysis allows rapid and reliable unmixing of fluorescence microscopy spectral images,” Optics express 20(12), 12729–12741 (2012).

49. E. Nehme, D. Freedman, R. Gordon, B. Ferdman, T. Michaeli, and Y. Shechtman, “Dense three dimensional localization microscopy by deep learning,” 1906.09957 [physics] (2019).

50. A. Speiser, S. C. Turaga, and J. H. Macke, “Teaching deep neural networks to localize sources in super-resolution microscopy by combining simulation-based learning and unsupervised learning,” 1907.00770 [cs, eess, stat] (2019).

51. A. Barsic, G. Grover, and R. Piestun, “Three-dimensional super-resolution and localization of dense clusters of single molecules,” Scientific reports 4, 5388 (2014).

